# Loss of microbial diversity and pathogen domination of the gut microbiota in critically ill patients

**DOI:** 10.1101/582494

**Authors:** Anuradha Ravi, Fenella D Halstead, Amy Bamford, Anna Casey, Nicholas M. Thomson, Willem van Schaik, Catherine Snelson, Robert Goulden, Ebenezer Foster-Nyarko, George M. Savva, Tony Whitehouse, Mark J. Pallen, Beryl A. Oppenheim

**Affiliations:** Quadram Institute Bioscience and University of East Anglia, Norwich, NR4 7UA, United Kingdom; NIHR Surgical Reconstruction and Microbiology Research Centre, Queen Elizabeth Hospital, Birmingham, B15 2GW, United Kingdom; Queen Elizabeth Hospital, University Hospitals Birmingham NHS Foundation Trust, Birmingham, B15 2GW, United Kingdom; Institute of Microbiology and Infection, University of Birmingham, Edgbaston, Birmingham B15 2TT, United Kingdom; McGill University, Montréal, QC H3G 2M1, Canada

**Author notes:** These authors contributed equally (Tony Whitehouse, Mark J. Pallen, Beryl A. Oppenheim).

**Keywords:** Intensive care unit, microbiome, gut microbiota, pathogens, shotgun metagenomics, antimicrobial resistance, critical illness, meropenem

## Abstract

**Background:** For long-stay patients on the adult intensive care unit, the gut microbiota plays a key role in determining the balance between health and disease. However, it remains unclear which ICU patients might benefit from interventions targeting the gut microbiota or the pathogens therein.

**Methods:** We undertook a prospective observational study of twenty-four ICU patients, in which serial faecal samples were subjected to shotgun metagenomic sequencing, phylogenetic profiling and microbial genome analyses.

**Results:** Two-thirds of patients experienced a marked drop in gut microbial diversity (to an inverse Simpson’s index of <4) at some stage during their stay in ICU, often accompanied by absence or loss of beneficial commensal bacteria. Intravenous administration of the broad-spectrum antimicrobial agent meropenem was significantly associated with loss of gut microbial diversity, but administration of other antibiotics, including piperacillin-tazobactam, failed to trigger statistically detectable changes in microbial diversity. In three quarters of ICU patients, we documented episodes of gut domination by pathogenic strains, with evidence of cryptic nosocomial transmission of *Enterococcus faecium*. In some patients we also saw domination of the gut microbiota by commensal organisms, such as *Methanobrevibacter smithii*.

**Conclusions:** Our results support a role for metagenomic surveillance of the gut microbiota and pave the way for patient-specific interventions that maintain or restore gut microbial diversity in the ICU.

## Background

For long-stay patients on the adult intensive care unit (ICU), as in other settings, the microbial community of the gut—the gut microbiota—plays a key role in determining the balance between health and disease [1–3]. Unfortunately, many life-saving measures applied to ICU patients can have negative impacts on the gut microbiota—examples include assisted ventilation, enteric feeds and a range of medications, including broad-spectrum antibiotics, proton-pump inhibitors, inotropes and opioids [4–6]. In recent years, interest has grown in protecting or restoring the integrity of the gut microbiome in ICU patients, using ecological approaches such as probiotics or faecal microbiota transplants [7–18]. Similarly, surveillance of pathogens and of antimicrobial resistance in the gut of critically ill patients has potential benefit in predicting infection and guiding treatment or infection control measures [19–21]. However, in the absence of high-precision approaches to the surveillance of complex microbial communities, it remains unclear which ICU patients might benefit from interventions affecting the gut microbiota and how such interventions should be targeted for optimum effect.

Fortunately, recent advances in sequencing and bioinformatics have made shotgun metagenomics an attractive approach in precision medicine [22, 23]. We therefore undertook a prospective observational study of twenty-four ICU patients, in which serial faecal samples were subjected to shotgun metagenomic sequencing, phylogenetic profiling and microbial genome analyses, with the aims of evaluating the utility of shotgun metagenomics in long-stay ICU patients, documenting the dynamics of the gut microbiota in this context and determining how it is affected by relevant clinical and demographic factors.

## Methods

### Study design and human subjects

Queen Elizabeth Hospital Birmingham is a university teaching hospital serving a population of approximately 1.5 million with a wide range of tertiary services including solid organ and bone marrow transplantation. Patients were enrolled for study participation if they were aged over 18 years, had been admitted to the ICU within the last 72 hours and were expected to remain there for more than 48 hours. Patients were considered evaluable if their first stool sample and at least one subsequent sample were collected on the ICU.

Patient information was collected on a case report form, which included information on gender, age, reason for admission, severity of disease scores, length of hospital stay prior to ICU admission, current and previous antibiotic therapy, blood markers, details of nutrition, drugs and relevant clinical microbiology results. The study started in May 2017 and ended in February 2018, when data and specimen collection for the thirtieth participant had been completed.

### Sample collection, storage and DNA extraction

The first faecal sample passed each calendar day by each enrolled patient on the ICU was collected and sent to the research team. Stool samples were aliquotted and then frozen at −20°C as soon as possible after collection. They were then shipped frozen to the Quadram Institute in Norwich, where they were stored at −80°C. Faecal samples were destroyed at the end of the study. Around 0.1 to 0.2 mg of frozen faecal sample was used for DNA extraction. The extraction was carried out using FastDNA Spin kit for Soil (MP Biomedicals, California, USA) according to the manufacturer’s instructions, except that 100 μl rather than 50 μl of DES elution buffer was used in the final elution.

Samples from extra-intestinal sites were collected when indicated on clinical grounds and processed by the hospital’s clinical microbiology laboratory using standard diagnostic procedures.

### Shotgun metagenomic sequencing

The DNA concentration was normalised using Qubit 4 (Invitrogen, Thermo Fisher, MA, USA) and sequencing libraries were prepared using the Nextera XT kit (Illumina). The DNA fragmented, tagged, cleaned and normalized according to the manufacturer’s recommendations. The quality of final pooled library was evaluated using Agilent 2200 Tape Station (Agilent) and the concentration was measured using Qubit 4 (Invitrogen, Thermo Scientific, MA, USA). Libraries were sequenced in batches on a NextSeq 550 using a high-output flow cell delivering 150-bp paired-end reads. The libraries were sequenced to a sequencing depth of ~ 2 Gbp/ sample.

Reads from the sequencer were uploaded on to virtual machines provided by the MRC CLIMB (Cloud Infrastructure for Microbial Bioinformatics) project using BaseMount [24, 25]. Initially, the sequences were assessed for quality using FastQC (version 0.11.5) and SeqKit with the parameter ‘stats’ [26, 27]. Quality filtering was performed using Trimmomatic (version 0.35) with default parameters [28]. Trimmomatic’s Illuminaclip function was used to remove Illumina adapters. Human sequences were removed by mapping reads towards human genome, Hg19 using BowTie2 version 2.3.4.1 [46] for mapping and SAMtools [29] to view with parameters -f 12 -F 256 to extract unmapped sequences and BEDtools bamtofastq to convert BAM to FASTQ files. [30]. Then, these sequences were deposited in the Sequence Read Archive under reference SUB5204757.

### Taxonomic profiling and statistical analysis

Forward and reverse paired reads were merged for each sample and fed as input to MetaPhlAn2 v2.7.7, which was used for taxonomic assignment of reads in each sample [31]. Metaphlan2 output was merged using the python script merge_metaphlan_tables.py. A species-only abundance table was created using Text Wrangler v5.5.2. Species that occurred only once and species with a relative abundance below 1% in the whole dataset were discarded. This abundance data table (Additional File 1) was used for diversity analyses.

Alpha diversity was assessed using the inverse Simpsons index calculated from the MetaPhlAn2 output using the vegan package (version 2.5-4) in R (version 3.5.2) [32]. Linear mixed models were used to estimate the fixed effects on alpha diversity of time since ICU admission, antibiotic use, time to sample storage and health status measures by SOFA score, and age and sex of the patient. The *nlme* package (version 3.1-137) in R (version 3.5.2) was used to estimate all models [32, 33].

Use of meropenem and piperacillin-tazobactam was coded individually because of their clinical importance and high use in our dataset, while all other antimicrobials were grouped together in a single variable ‘other antimicrobials’ for the final multivariable analysis. To account for long-term effects of antibiotics on microbial diversity and absence of data on when antibiotics were started, antibiotic use variables were coded at each sampling point into one of four levels: no use, starting use, ongoing use and historic use. Episodes were classified as ‘starting’ if the antibiotic was started on same day the sample was taken; ‘ongoing’ if the antibiotic was still being administered on the day of a sample being taken; ‘historic’ if the antibiotic had been used prior to the date of sample collection but was no longer being administered. Data from 228 samples was included in the final analysis, with nine samples excluded because the SOFA score was missing. This dataset included 42 samples taken while meropenem was administered and 44 while piperacillin-tazobactam was administered. Complete-case analysis was used in the case of missing data. Random patient-level effects on intercept and slope (linear change in diversity over time) were included. An auto-regressive correlation structure (AR1) in discrete time was used to account for the residual autocorrelation evident from initial mixed models. Residual autocorrelation in the final model was minimal.

### Metagenome-assembled genomes

For metagenomic binning, reads from each patient was co-assembled into contigs using MEGAHIT v1.1.3. [34]. Next, anvi’o version 5.1 was used for mapping, binning, refining and visualising the bins. [35]. In brief, ‘anvi-gen-contigs-database’ was used with default settings to profile the contigs using Prodigal v2.6.3 and identify open reading frames [36]. Then, ‘anvi-run-hmms’ was used with default settings to identify bacterial, archaeal and fungal single copy gene collections using HMMER [37] and ‘anvi-run-cogs’ was used to predict gene functions in the contigs by using NCBI’s Cluster of Orthologous Groups database. Taxonomy of the contigs was predicted using Centrifuge v1.0.3-beta [38] and added to the database using ‘anvi-import-taxonomy-for-genes’ function. Reads from each sample of the respective patient was mapped to their corresponding co-assembled contigs using Bowtie v2.2.3 and converted into sorted and indexed bam files using Samtools v1.9. Then, ‘anvi-profile’ was used to profile each bam file to estimate coverage and detection statistics for every contig in each sample. Then, ‘anvi-merge’ is used to combine the profiles of each sample and create a merged anvi’o profile. Then, ‘anvi-interactive’ was used to interactively visualise the distribution of the bins and identify metagenome-assembled genomes (MAGs).

We classified a genome bin as a MAG if it was more than 80% complete and its redundancy was below 10%. Each bin was then refined using ‘anvi-refine’ based on tetranucleotide frequency, mean coverage, completion and redundancy. The program ‘anvi-summarize’ was used to generate a HTML output stat and FASTA file with the recovered MAGs. To reconfirm the completion and redundancy of the MAGs, CheckM v1.0.13 was used [39]. To recover MAGs for the fungal genomes, ‘anvi-run-hmms’ was used with BUSCO [40], a collection of 83 eukaryotic single copy genes, ‘anvi-compute-completeness’ was used to identify completion and ‘anvi-interactive’ was used to recover the MAGS.

For low-abundance pathogens that had been identified by MetaPhlAn2 but could not be recovered using Anvi’o, we constructed sets of completed taxon-specific reference genomes for each potential pathogen. Reference sequences were downloaded using the ncbi-genome-download script [45]. We then mapped the metagenome from each sample against the relevant reference dataset using BowTie2 version 2.3.4.1 [46]. The mapped reads were recovered using BEDtools bamtofastq and assembled into contigs using SPAdes (version 3.11.1) [49] and annotated using Prokka (version 1.12) [41]. Completion and contamination of these MAGs were assessed using CheckM. The coverage of the resulting draft genome sequences was calculated after mapping reads back to the assemblies using BowTie2 and visualised with Qualimap2 [42]. To confirm species identity, average nucleotide identity was calculated from BLAST searches [43] or by using the online ANI/AAI matrix tool [44].

Resistance genes in the MAGs were detected using ABRicate v0.8.10 with ResFinder and CARD databases. [45]. The reports from the individual samples was compiled using the ‘— summary’ option. Antifungal resistance genes were manually identified in the annotated MAGs and curated using the Candida Genome Database. [46]. SNP distance matrices between the MAGs were calculated using Snippy v3.1 incorporating Freebayes v1.1.0 for SNP detection [47].

### Pathogen culture

For the isolation of *Escherichia coli, Candida albicans* and *Enterococcus faecium*, two separate aliquots (0.1 – 0.2 g) of each stool sample were loaded into 1.5ml microcentrifuge tubes under aseptic conditions. 1ml of physiological saline (0.85%) was added and the saline-stool samples were vortexed for two minutes at maximum speed to homogenise the samples completely. The homogenised samples were taken through eight 10-fold serial dilutions and 100 µl aliquots from each dilution were dispensed on to Tryptone-Bile-X-Glucoronide agar, Sabauraud-Dextrose Agar and Slanetz and Bartley medium (Oxoid). Both aliquots were plated in triplicate. The sample suspensions were spread on the plates using the cross-hatching method for confluent growth. Inoculated plates incubated at 37°C for 18 – 24 hours (for Tryptone-Bile-X-Glucoronide agar, Sabauraud-Dextrose Agar) or for 48 hours on Slanetz and Bartley medium.

Following incubation, the plates were examined for growth. On Tryptone-Bile-X-Glucoronide agar, raised blue-green colonies with entire margins were taken as indicative of the growth of *E. coli*. On Sabauraud-Dextrose Agar, raised white-to-cream entire colonies with yeast-like appearance were scored as *Candida.* On Slanetz and Bartley medium, smooth pink-to-red colonies with a whitish margin were indicative of the growth of *Enterococcus*. Colonies were counted on the dilution plate that showed the highest number of discrete colonies and the colony count for each of the triplicate plates per dilution was recorded.

## Results

We initially recruited thirty serially recruited adult patients who were expected to stay on the ICU for >48 hours. As is typical of ICU patients, the study population was heterogeneous, including patients with little or no previous medical history (e.g. suffering from trauma or intracranial haemorrhage) as well as individuals with complex and chronic clinical conditions and varying immune function (Additional File 2). A total of 236 faecal samples were collected, with a median of three days between samples from each patient (Additional File 3). A set of twenty-four long-stay ICU patients who provided more than five samples was selected for further study (Table 1).

**Table 1.**
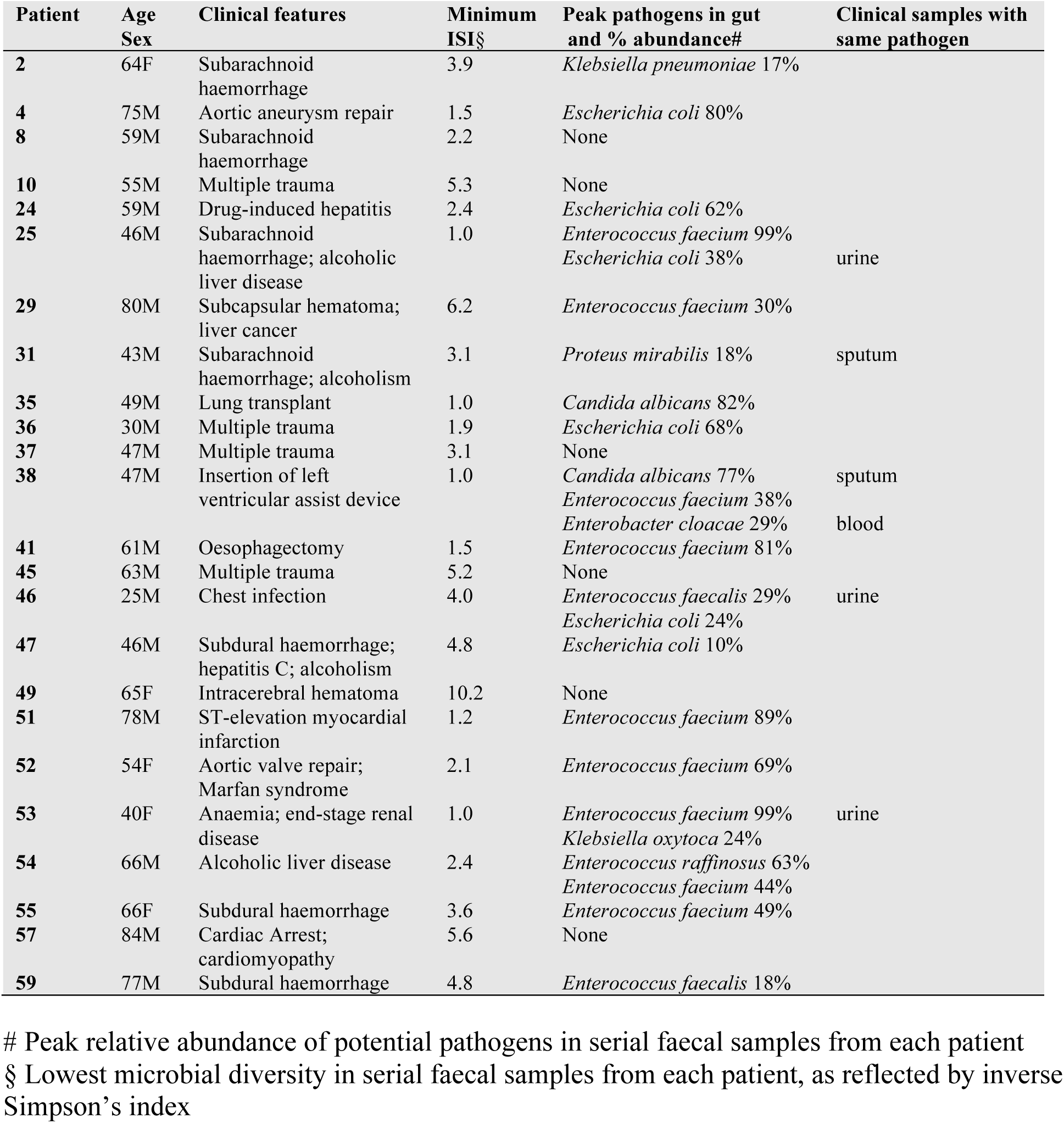
Clinical features and gut microbial ecology of ICU patients

To track the gut microbial dynamics of individual patients, we performed metagenomic sequencing of serial faecal samples, followed by community analysis to determine the relative abundance of microbial species and to assess microbial diversity using the inverse Simpson’s index (Additional Files 3, 4). The median time to receipt of a sample (where timings were available) was 2.6 hours: 70% of samples were received within 6 hours and 87% within 12 hours. We found no association between microbial diversity and time to receipt of sample.

### Loss of gut microbial diversity with meropenem

We found no general trend towards decreased gut microbial diversity with time spent in ICU. Similarly, we found no statistically significant associations between microbial diversity and stool consistency or Sequential Organ Failure Assessment (SOFA) score, which reflects overall health (Table 2). However, in 11 of 24 patients, microbial diversity in the final sample was lower than in the initial sample (Table 2) and in two-thirds of patients, we saw a fall in diversity at some stage during their stay in ICU to an inverse Simpson’s index of <4: a figure which has been associated with decreased survival in immunocompromised patients [48] (Table 1; Additional File 4). Remarkably, in a third of our patients, diversity fell, in at least one sample, to a precipitously low level, with an inverse Simpson’s index < 2, echoing findings from previous studies that used 16S amplicon sequencing [49, 50].

**Table 2.** Gut Microbial Diversity and Clinical factors. Coefficients from a mixed effects regression model measuring the association between faecal microbial alpha diversity (inverse Simpson’s index) and demographics and clinical factors. Total N=228 samples included in the analysis.

|  | Unit/level | N / Mean (SD) | Coefficient | 95% Confidence interval | p-value |
| --- | --- | --- | --- | --- | --- |
| Age | per year | 54.6 (14.8) | 0.03 | (-0.03, 0.09) | 0.382 |
| Sex | male vs. female | 170 | -0.14 | (-2.40, 2.12) | 0.897 |
| Time since admission | per day | 18.1 (12.5) | -0.03 | (-0.10, 0.04) | 0.421 |
| Meropenem | No use | 114 | 0 | Reference |  |
|  | Ongoing | 42 | -1.82 | (-3.40, -0.25) | **0.024\*** |
|  | Starting | 7 | -1.30 | (-3.03, 0.44) | 0.143 |
|  | History | 65 | -1.29 | (-2.92, 0.35) | 0.122 |
| Piperacillin-tazobactam | No use | 51 | 0 | Reference |  |
|  | Ongoing | 44 | 0.66 | (-1.09, 2.42) | 0.456 |
|  | Starting | 4 | 1.50 | (-0.87, 3.87) | 0.214 |
|  | History | 129 | 0.83 | (-0.92, 2.58) | 0.350 |
| Other antimicrobial | No use | 32 | 0 | Reference |  |
|  | Ongoing | 55 | -1.16 | (-3.12, 0.79) | 0.242 |
|  | Starting | 8 | -0.03 | (-2.15, 2.09) | 0.980 |
|  | History | 133 | -0.99 | (-2.83, 0.85) | 0.290 |
| Bristol Stool Index | 1-3 | 9 | 0 | Reference |  |
|  | 4 | 28 | -0.54 | (-2.25, 1.17) | 0.536 |
|  | 5 | 48 | 0.32 | (-1.40, 2.04) | 0.715 |
|  | 6 | 75 | -0.19 | (-1.83, 1.46) | 0.823 |
|  | 7 | 62 | -0.70 | (-2.41, 1.02) | 0.423 |
|  | Missing | 6 | 0.04 | (-2.22, 2.30) | 0.975 |
| SOFA score | per point | 6.1 (3.4) | -0.15 | (-0.31, 0.01) | 0.065 |

SOFA score: sequential organ failure score, higher values = greater morbidity
\*: p<0.05
N: number of samples with each level of covariate
SD: standard deviation

All but one of our ICU patients were treated with antimicrobial chemotherapy, varying from one to six classes of antibacterial or antifungal agent. Surprisingly, in these patients, administration of most antibiotics, including the broad-spectrum agent piperacillin-tazobactam, failed to trigger statistically detectable changes in microbial diversity, despite apparent sensitivity of gut commensals to such agents [6]. However, current use of the intravenous agent meropenem was significantly associated with loss of gut microbial diversity in our ICU patients, confirming similar findings with healthy adults (change in inverse Simpsons index −1.8, 95% CI=−3.4 to −0.25; p=0.024; Table 2) [51].

### Loss of beneficial microorganisms

For many patients during their stay in ICU, we observed an absence or drastic reduction in the gut of microorganisms thought to confer beneficial effects on the host. Loss of beneficial organisms was particularly marked after administration of meropenem (Figure 1). Examples include *Faecalibacterium prauznetzii*, a butyrate-producing bacterium with anti-inflammatory properties; *Akkermansia muciniphila*, a mucin-degrading bacterium inversely related to diabetes and inflammation (both undetectable in final samples from 18 patients) [52, 53]; *Bacteroides thetaiotaomicron*, which inhibits colonisation of the gut by *Candida albicans* (undetectable in all samples from eight patients; lost in four) [54]; *Clostridium bolteae* (lost in seven patients) and *Blautia producta* (undetectable in 13 patients), which strengthen colonisation resistance against vancomycin-resistant *Enterococci* [55]; and candidate probiotic species such as *Bifidobacterium longum* (undetectable in all samples from ten patients; lost in six); and *Bifidobacterium adolescentis* (undetectable in all samples from nineteen patients; lost in four) [56].

**Figure 1.**
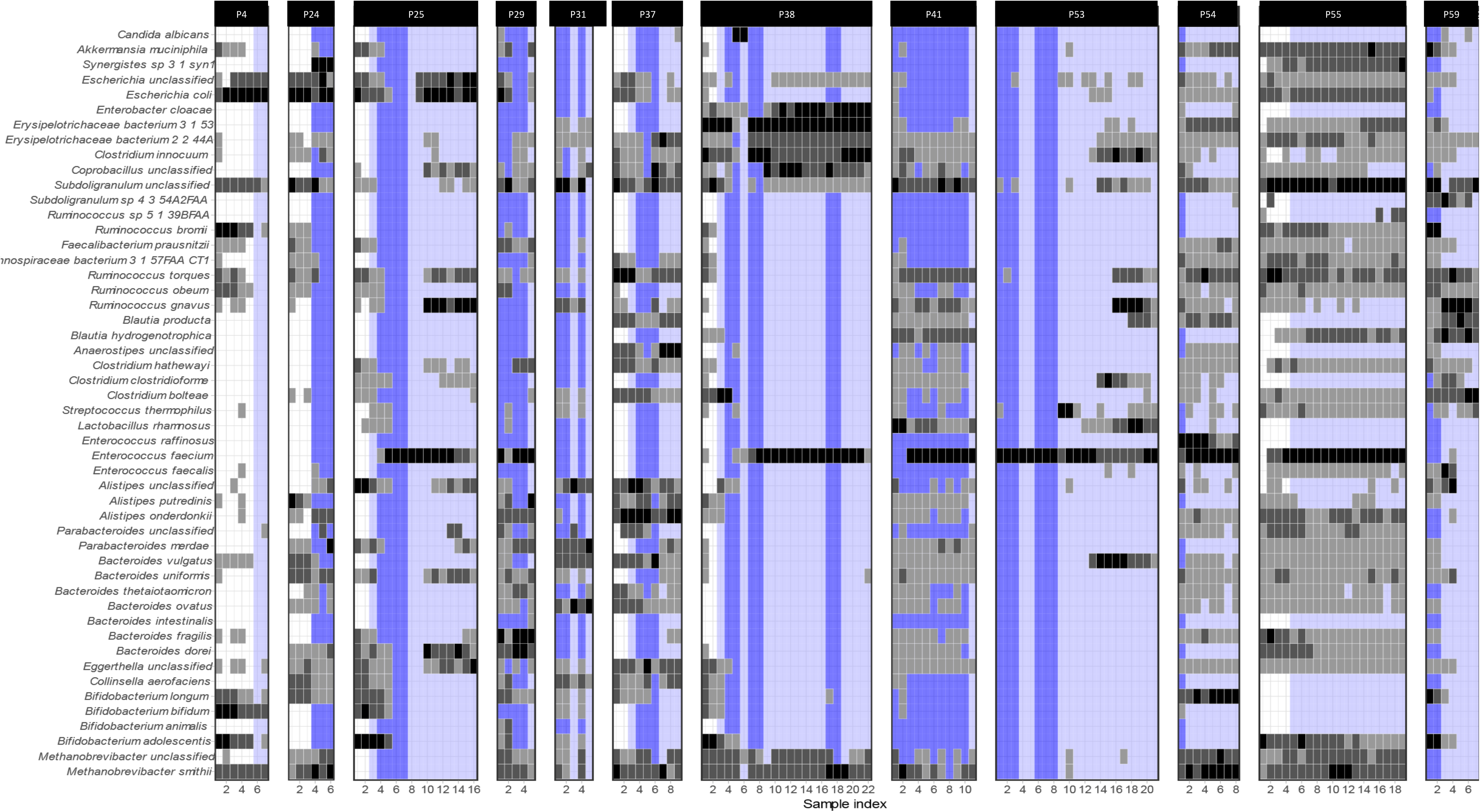
Relative abundance of gut microorganisms among patients who began meropenem during the study. This heat map shows the top 50 taxa by average relative abundance across the whole dataset. Grey-scale shading of cells shows relative abundance: 0 (no shading) 0-1% (light grey), 1-10% (mid-grey) and >10% (dark grey). Coloured shading of columns reflects meropenem use: no use (blank); ongoing use (dark blue); starting meropenem or a history of meropenem (light blue)

### Domination of the gut by individual pathogens and commensals

In 75% of the long-stay ICU patients, we saw marked increases in the relative abundance of individual pathogens in stool samples. These included ESKAPE pathogens (*Enterococcus faecium, Klebsiella pneumoniae, Enterobacter cloacae*), other species of Enterobacteriaceae (*Escherichia coli, Klebsiella oxytoca, Proteus mirabilis*) and enterococci (*E. faecalis* and *E. raffinosus*) and the fungal pathogen *Candida albicans*. During these episodes of pathogen domination, the relative abundance of the pathogen often exceeded 50% of sequence reads—in one patient, patient 53, in seven consecutive samples, >80% of evaluable sequences were assigned to *E. faecium* (Figure 2).

**Figure 2.**
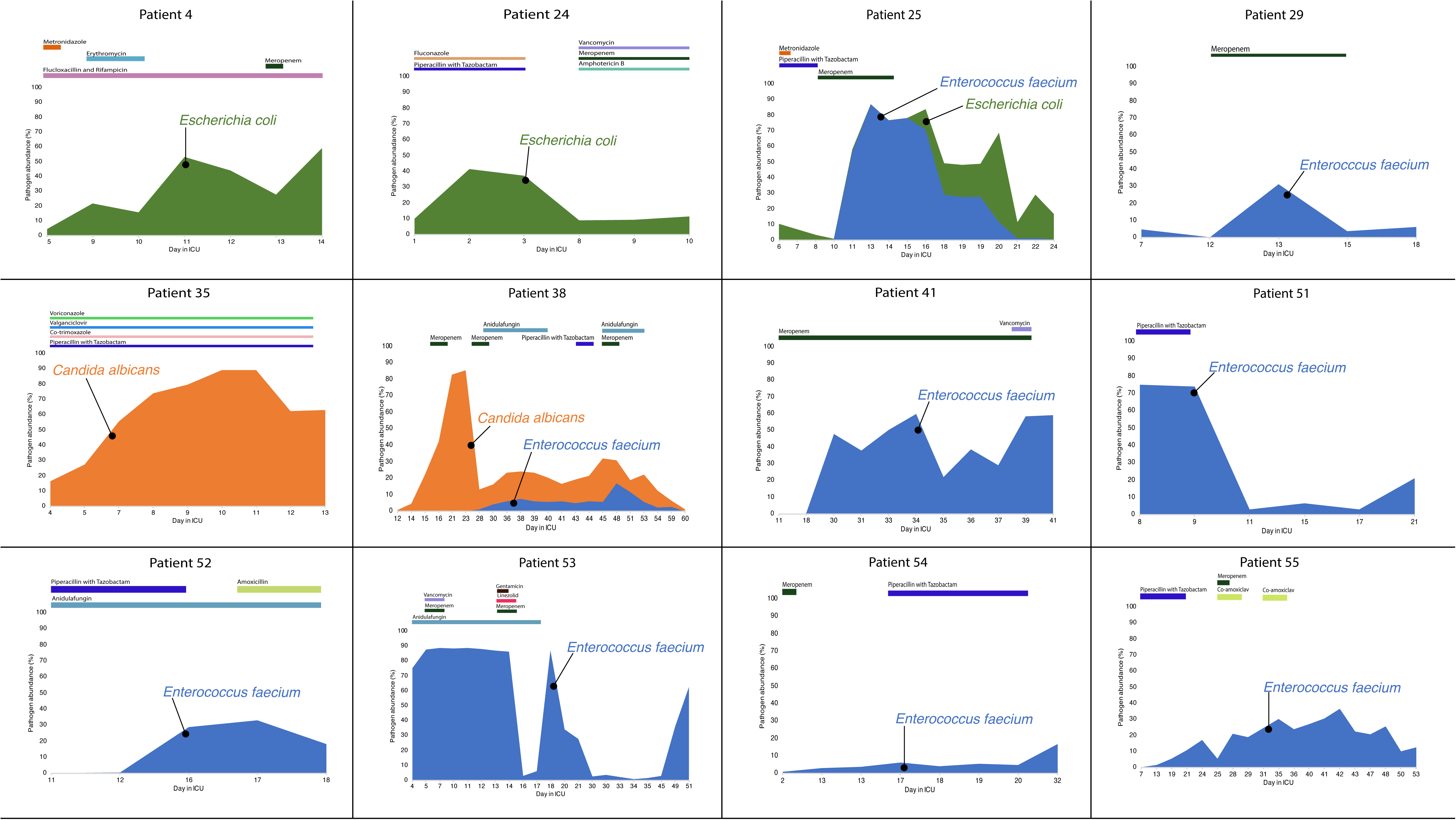
Pathogen domination of the gut microbiota. Timelines for patients showing pathogen domination, with relative abundance assessed by percentage of reads mapping to metagenome-assembled genomes. Various antibiotics were given for treatment purposes during the study period.

Through quantitative culture of pathogens from serial faecal samples, we documented changes in absolute abundance in some cases (Additional File 5). In six patients, the same species of pathogen was isolated from clinical samples from outside the gut (Table 1).

Antibiotics are known to provoke overgrowth in the gut of microbial species not known to be pathogens [51, 57, 58]. We saw the relative abundance of the archaeon *Methanobrevibacter smithii* exceed 10% of reads in nine ICU patients—in one sample, this organism accounted for 50% of sequence reads. Other apparent commensals showing rises in relative abundance to >50% include *Streptococcus thermophilus, Alistipes onderdonkii, Bifidobacterium longum*, an unnamed species from the *Erysipelotrichaceae, Ruminococcus torques* and an unclassified species of *Subdoligranulum* (Figure 1; Additional File 1).

### Cryptic nosocomial transmission

Through a combination of co-assembly, *de novo* binning of metagenomic reads and evaluation of single copy bacterial and fungal core genes, we obtained metagenome-assembled genomes (MAGs) of potential pathogens and used them to reconstruct pathogen biology and epidemiology, including multi-locus sequence types (Additional File 6). We found that pathogen blooms within an individual patient were typically clonal, i.e. caused by a single strain, although there was often a cloud of diversity in single-nucleotide polymorphisms among genomes from multiple samples (Additional File 7). Pathogens dominating the gut microbiota in ICU patients were also typically inherently resistant to antibiotics (*Candida albicans*) or possessed genes associated with antimicrobial resistance—vancomycin-resistance genes were detected in two strains of *E. faecium* and aminoglycoside resistance genes in two strains of *E. coli*, one of which also encoded an extended-spectrum beta-lactamase (Additional File 8).

In one patient, Patient 25, where we sequenced multiple isolates cultured from the first sample from the patient, we detected two strains of *E. coli* belonging to distinct sequence types, ST131 and ST315. However, the ST131 strain was subsequently lost after administration of antibiotics. Enterococcal blooms were seen in eleven patients (Figure 2). In six cases, the dominant strain belonged not just to the same species, *E. faecium*, but also to the same sequence type, ST80, which is a well-documented cause of nosocomial outbreaks across the globe [59–61]. We found that the *E. faecium* MAGs belonging to ST80 in patients 51, 54, 55 differed by less than twenty SNPs (Additional File 8), providing strong evidence of cross-colonization with a common strain, spreading between patients and/or from a common source in the hospital. Interestingly, all three patients had overlapping stays in adjacent rooms on the ICU.

## Discussion

Here, we have shown the utility of shotgun metagenomics in in ICU patients in surveillance of the gut microbiota, documenting the loss of gut microbial diversity and domination of the gut by drug-resistant pathogens. Our use of shotgun metagenomics confirms results of previous studies on ICU patients using less powerful sequence-based approaches, linking loss of gut microbial diversity to adverse clinical outcomes and loss of colonization resistance, [19–21, 49, 50, 62–69]. However, with shotgun metagenomics, we have been able to reconstruct informative metagenome-assembled genomes, allowing us to characterise pathogens, identify resistance determinants and document cryptic nosocomial transmission of a clone of *Enterococcus faecium* that colonised three patients.

It is well established that administration of antibiotics leads to loss of diversity in the gut microbiota [70–73]. Nonetheless, although all but one of our patients received antibiotics, we saw a statistically significant loss of diversity—and marked loss of beneficial organisms—only after administration of meropenem (Figure 1). Similar profound and longstanding effects of this agent on gut microbial diversity have been documented in healthy adults [51]. Although we were unable to detect any effect of other antimicrobials, given the small sample size, we cannot rule out small but significant effects for less commonly used agents.

The contrast between the effect of meropenem and apparent lack of effect of other broad-spectrum agents such as piperacillin-tazobactam suggests that pharmacokinetics plays a key role in determining impact on the gut microbiota and that there is scope for tailoring antibiotic regimes to spare the gut microbiota, building on previous studies confirming the low-risk status of ureidopenicillins such as piperacillin on the risk of *Clostridioides difficile* infection or colonisation with vancomycin-resistant enterococci [74, 75].

We have used shotgun metagenomics to document domination of the gut microbiota by microbial pathogens in most ICU patients. Although from sequences alone, it is hard to determine whether increases in relative abundance of pathogens reflect an increase in the biomass of pathogens or simply a loss of commensals [76, 77], we were able to use microbial culture to confirm that, at least in some cases, there was a genuine increase in the absolute abundance of the pathogen.

In a quarter of our patients, in line with other similar attempts at sequence-based surveillance in vulnerable patients[19, 20], the same species of pathogen was isolated from clinical samples from outside the gut. However, as our clinical isolates were not subjected to genome sequencing, we cannot be certain that they belonged to the strains associated with domination of the gut.

Interesting, we also saw episodes of ecological domination by apparent commensals. The significance of these episodes remains uncertain. A recent study has suggested that commensal bacteria carry diverse uncharacterised resistance genes that contribute to their selection after antibiotic therapy [78]. It is worth noting that *M. smithii*, like other Archaea, is intrinsically resistant to antibiotics as a result of its distinctive non-bacterial biology [79].

## Conclusions

Here, we have shown that surveillance of the gut microbiota in long-stay ICU patients using shotgun metagenomics is capable of detecting episodes of low diversity and pathogen domination, as well as providing genome-level resolution of colonising pathogens and evidence of cryptic nosocomial transmission. We have also shown that use of meropenem is associated with ecological disruption of the gut microbiota. These observations pave the way for precise patient-specific interventions that protect the gut microbiota (e.g. enhanced infection control, tailored use of microbiota-sparing antibiotics, oral administration of antibiotic-absorbing charcoal or of a beta-lactamase) [80, 81].

Although we failed to find a link between gut microbial diversity or pathogen domination of the gut and clinical outcomes in our group of ICU patients, such evidence has been documented for similar groups of vulnerable patients [19, 21, 63], where ecological approaches to restoring gut microbial diversity, such as faecal microbiota transplants, are under evaluation [16–18, 82, 83]. Similar intervention studies—underpinned by the kind of metagenomic surveillance we have established here—are likely to clarify whether maintenance or restoration of gut microbial diversity influences clinical outcomes in long-stay ICU patients.

## Abbreviations

ICU: Intensive Care Unit
MAG: metagenome-assembled genome
SOFA: Sequential Organ Failure Assessment

## Declarations

### Ethics approval and consent to participate

Wales Research Ethics Committee 3 granted ethical approval for the study under the auspices of the UK’s National Research Ethics Service (Reference number 17/WA/0073; Integrated Research Application System ID 222006) and the study was conducted according to the World Medical Association’s Declaration of Helsinki. Informed consent was obtained from the patient or from the patient’s consultee (surrogate decision-maker) using standard consent procedures for clinical studies of ICU populations within the UK, which include provision for patients who lack capacity.

### Consent for publication

Not applicable

### Availability of data and material

Metagenome sequences have been deposited in the Sequence Read Archive (https://www.ncbi.nlm.nih.gov/sra) under Bioproject reference SUB5204757

### Authors’ contributions

BAO, TW, RG, FDH and AC designed and supervised the clinical study. FDH, RG, AC, AB and CS carried out the clinical components, sample and data collection. AR, GS, EF-N, WvS. and MJP designed and/or carried out sequencing, computational cultural or statistical analysis. MJP wrote the paper. All authors read and approved the paper.

## Acknowledgements

We thank ICU staff for their tireless work and contributions to this study.

## Additional Files

**Additional File 1.** Stool data and MetaPhlAn2 output

**Additional File 2.** Patient data

**Additional File 3.** Sequence data

**Additional File 4.** Diversity indices

**Additional File 5.** Pathogen abundance on culture

**Additional File 6.** Metagenome-assembled genomes statistics

**Additional File 7.** Identification of antibiotic resistance genes

**Additional File 8.** SNP matrices for enterococcal metagenome-assembled genomes

## References

1. Kim S, Covington A, Pamer EG: The intestinal microbiota: Antibiotics, colonization resistance, and enteric pathogens. Immunol Rev 2017, 279:90–105.

2. Feng Q, Chen WD, Wang YD: Gut microbiota: An integral moderator in health and disease. Front Microbiol 2018, 9:151.

3. Dickson RP: The microbiome and critical illness. Lancet Respir Med 2016, 4:59–72.

4. Chang SJ, Huang HH: Diarrhea in enterally fed patients: blame the diet. Curr Opin Clin Nutr Metab Care 2013, 16:588–594.

5. Lustri BC, Sperandio V, Moreira CG: Bacterial chat: Intestinal metabolites and signals in host-microbiota-pathogen interactions. Infect Immun 2017, 85

6. Maier L, Pruteanu M, Kuhn M, Zeller G, Telzerow A, Anderson EE, Brochado AR, Fernandez KC, Dose H, Mori H, Patil KR, Bork P, Typas A: Extensive impact of non-antibiotic drugs on human gut bacteria. Nature 2018, 555:623–628.

7. Wischmeyer PE, McDonald D, Knight R: Role of the microbiome, probiotics, and ‘dysbiosis therapy’ in critical illness. Curr Opin Crit Care 2016, 22:347–353.

8. Pamer EG: Resurrecting the intestinal microbiota to combat antibiotic-resistant pathogens. Science 2016, 352:535–538.

9. Manges AR, Steiner TS, Wright AJ: Fecal microbiota transplantation for the intestinal decolonization of extensively antimicrobial-resistant opportunistic pathogens: a review. Infect Dis (Lond) 2016, 48:587–592.

10. Morrow LE, Wischmeyer P: Blurred lines: Dysbiosis and probiotics in the icu. Chest 2017, 151:492–499.

11. Haak BW, Levi M, Wiersinga WJ: Microbiota-targeted therapies on the intensive care unit. Curr Opin Crit Care 2017, 23:167–174.

12. Wolff NS, Hugenholtz F, Wiersinga WJ: The emerging role of the microbiota in the ICU. Crit Care 2018, 22:78.

13. McClave SA, Patel J, Bhutiani N: Should fecal microbial transplantation be used in the ICU. Current Opinion in Critical Care 2018, 24:105–111.

14. Limketkai BN, Hendler S, Ting PS, Parian AM: Fecal microbiota transplantation for the critically ill patient. Nutr Clin Pract 2018,

15. Ruppe E, Martin-Loeches I, Rouze A, Levast B, Ferry T, Timsit JF: What’s new in restoring the gut microbiota in ICU patients? Potential role of faecal microbiota transplantation. Clin Microbiol Infect 2018, 24:803–805.

16. Dinh A, Fessi H, Duran C, Batista R, Michelon H, Bouchand F, Lepeule R, Vittecoq D, Escaut L, Sobhani I, Lawrence C, Chast F, Ronco P, Davido B: Clearance of carbapenem-resistant Enterobacteriaceae vs vancomycin-resistant enterococci carriage after faecal microbiota transplant: a prospective comparative study. J Hosp Infect 2018, 99:481–486.

17. Davido B, Batista R, Fessi H, Michelon H, Escaut L, Lawrence C, Denis M, Perronne C, Salomon J, Dinh A: Fecal microbiota transplantation to eradicate vancomycin-resistant enterococci colonization in case of an outbreak. Med Mal Infect 2018,

18. Suez J, Zmora N, Zilberman-Schapira G, Mor U, Dori-Bachash M, Bashiardes S, Zur M, Regev-Lehavi D, Ben-Zeev Brik R, Federici S, Horn M, Cohen Y, Moor AE, Zeevi D, Korem T, Kotler E, Harmelin A, Itzkovitz S, Maharshak N, Shibolet O, Pevsner-Fischer M, Shapiro H, Sharon I, Halpern Z, Segal E, Elinav E: Post-antibiotic gut mucosal microbiome reconstitution is impaired by probiotics and improved by autologous FMT. Cell 2018, 174:1406–1423.e16.

19. Taur Y, Xavier JB, Lipuma L, Ubeda C, Goldberg J, Gobourne A, Lee YJ, Dubin KA, Socci ND, Viale A, Perales MA, Jenq RR, van den Brink MR, Pamer EG: Intestinal domination and the risk of bacteremia in patients undergoing allogeneic hematopoietic stem cell transplantation. Clin Infect Dis 2012, 55:905–914.

20. Tamburini FB, Andermann TM, Tkachenko E, Senchyna F, Banaei N, Bhatt AS: Precision identification of diverse bloodstream pathogens in the gut microbiome. Nat Med 2018, 24:1809–1814.

21. Freedberg DE, Zhou MJ, Cohen ME, Annavajhala MK, Khan S, Moscoso DI, Brooks C, Whittier S, Chong DH, Uhlemann AC, Abrams JA: Pathogen colonization of the gastrointestinal microbiome at intensive care unit admission and risk for subsequent death or infection. Intensive Care Med 2018,

22. Pallen MJ: Diagnostic metagenomics: potential applications to bacterial, viral and parasitic infections. Parasitology 2014, 141:1856–1862.

23. Hillmann B, Al-Ghalith GA, Shields-Cutler RR, Zhu Q, Gohl DM, Beckman KB, Knight R, Knights D: Evaluating the information content of shallow shotgun metagenomics. mSystems 2018, 3

24. Connor TR, Loman NJ, Thompson S, Smith A, Southgate J, Poplawski R, Bull MJ, Richardson E, Ismail M, Thompson SE, Kitchen C, Guest M, Bakke M, Sheppard SK, Pallen MJ: CLIMB (the Cloud Infrastructure for Microbial Bioinformatics): an online resource for the medical microbiology community. Microb Genom 2016, 2:e000086.

25. BaseMount: A Linux command line interface for BaseSpace [https://blog.basespace.illumina.com/2015/07/23/basemount-a-linux-command-line-interface-for-basespace/]

26. FastQC: a quality control tool for high throughput sequence data. [http://www.bioinformatics.babraham.ac.uk/projects/fastqc]

27. Shen W, Le S, Li Y, Hu F: SeqKit: A Cross-Platform and Ultrafast Toolkit for FASTA/Q File Manipulation. PLoS One 2016, 11:e0163962.

28. Bolger AM, Lohse M, Usadel B: Trimmomatic: a flexible trimmer for Illumina sequence data. Bioinformatics 2014, 30:2114–2120.

29. Li H, Handsaker B, Wysoker A, Fennell T, Ruan J, Homer N, Marth G, Abecasis G, Durbin R, 1000 GPDPS: The Sequence Alignment/Map format and SAMtools. Bioinformatics 2009, 25:2078–2079.

30. Quinlan AR: BEDTools: The Swiss-Army Tool for Genome Feature Analysis. Curr Protoc Bioinformatics 2014, 47:11.12.1-34.

31. Truong DT, Franzosa EA, Tickle TL, Scholz M, Weingart G, Pasolli E, Tett A, Huttenhower C, Segata N: MetaPhlAn2 for enhanced metagenomic taxonomic profiling. Nat Methods 2015, 12:902–903.

32. R-Core-Team.: R: A language and environment for statistical computing. R Foundation for Statistical Computing, Vienna, Austria. 2018,

33. nlme: Linear and Nonlinear Mixed Effects Models. R package version 3.1-137 [https://CRAN.R-project.org/package=nlme]

34. Li D, Luo R, Liu CM, Leung CM, Ting HF, Sadakane K, Yamashita H, Lam TW: MEGAHIT v1.0: A fast and scalable metagenome assembler driven by advanced methodologies and community practices. Methods 2016, 102:3–11.

35. Eren AM, Esen ÖC, Quince C, Vineis JH, Morrison HG, Sogin ML, Delmont TO: Anvi’o: an advanced analysis and visualization platform for ‘omics data. PeerJ 2015, 3:e1319.

36. Hyatt D, Chen GL, Locascio PF, Land ML, Larimer FW, Hauser LJ: Prodigal: prokaryotic gene recognition and translation initiation site identification. BMC Bioinformatics 2010, 11:119.

37. Potter SC, Luciani A, Eddy SR, Park Y, Lopez R, Finn RD: HMMER web server: 2018 update. Nucleic Acids Res 2018, 46:W200–W204.

38. Kim D, Song L, Breitwieser FP, Salzberg SL: Centrifuge: rapid and sensitive classification of metagenomic sequences. Genome Res 2016, 26:1721–1729.

39. Parks DH, Imelfort M, Skennerton CT, Hugenholtz P, Tyson GW: CheckM: assessing the quality of microbial genomes recovered from isolates, single cells, and metagenomes. Genome Res 2015, 25:1043–1055.

40. Waterhouse RM, Seppey M, Simão FA, Manni M, Ioannidis P, Klioutchnikov G, Kriventseva EV, Zdobnov EM: BUSCO applications from quality assessments to gene prediction and phylogenomics. Mol Biol Evol 2017,

41. Seemann T: Prokka: rapid prokaryotic genome annotation. Bioinformatics 2014, 30:2068–2069.

42. Okonechnikov K, Conesa A, García-Alcalde F: Qualimap 2: advanced multi-sample quality control for high-throughput sequencing data. Bioinformatics 2016, 32:292–294.

43. Altschul SF, Gish W, Miller W, Myers EW, Lipman DJ: Basic local alignment search tool. J Mol Biol 1990, 215:403–410.

44. Rodriguez-R LM, Konstantinidis KT: The enveomics collection: a toolbox for specialized analyses of microbial genomes and metagenomes. PeerJ Preprints 2016, 4:e1900v1.

45. ABRicate [https://github.com/tseemann/abricate]

46. Skrzypek MS, Binkley J, Binkley G, Miyasato SR, Simison M, Sherlock G: The Candida Genome Database (CGD): incorporation of Assembly 22, systematic identifiers and visualization of high throughput sequencing data. Nucleic Acids Res 2017, 45:D592–D596.

47. Snippy: rapid haploid variant calling and core SNP phylogeny [https://github.com/tseemann/snippy]

48. Pamer EG, Taur Y, Jenq R, van den Brink MRM: Impact of the intestinal microbiota on infections and survival following hematopoietic stem cell transplantation. Blood 2014, 124:SCI–48.

49. Zaborin A, Smith D, Garfield K, Quensen J, Shakhsheer B, Kade M, Tirrell M, Tiedje J, Gilbert JA, Zaborina O, Alverdy JC: Membership and behavior of ultra-low-diversity pathogen communities present in the gut of humans during prolonged critical illness. Mbio 2014, 5:e01361–14.

50. McDonald D, Ackermann G, Khailova L, Baird C, Heyland D, Kozar R, Lemieux M, Derenski K, King J, Vis-Kampen C, Knight R, Wischmeyer PE: Extreme dysbiosis of the microbiome in critical illness. mSphere 2016, 1:207.

51. Palleja A, Mikkelsen KH, Forslund SK, Kashani A, Allin KH, Nielsen T, Hansen TH, Liang S, Feng Q, Zhang C, Pyl PT, Coelho LP, Yang H, Wang J, Typas A, Nielsen MF, Nielsen HB, Bork P, Wang J, Vilsbøll T, Hansen T, Knop FK, Arumugam M, Pedersen O: Recovery of gut microbiota of healthy adults following antibiotic exposure. Nat Microbiol 2018, 3:1255–1265.

52. Martín R, Bermúdez-Humarán LG, Langella P: Searching for the bacterial effector: The example of the multi-skilled commensal bacterium Faecalibacterium prausnitzii. Front Microbiol 2018, 9:1298.

53. Cani PD, de Vos WM: Next-Generation Beneficial Microbes: The Case of Akkermansia muciniphila. Front Microbiol 2017, 8:1765.

54. Fan D, Coughlin LA, Neubauer MM, Kim J, Kim MS, Zhan X, Simms-Waldrip TR, Xie Y, Hooper LV, Koh AY: Activation of HIF-1α and LL-37 by commensal bacteria inhibits Candida albicans colonization. Nat Med 2015, 21:808–814.

55. Caballero S, Kim S, Carter RA, Leiner IM, Sušac B, Miller L, Kim GJ, Ling L, Pamer EG: Cooperating commensals restore colonization resistance to vancomycin-resistant enterococcus faecium. Cell Host & Microbe 2017, 21:592–602.e4.

56. Arboleya S, Watkins C, Stanton C, Ross RP: Gut bifidobacteria populations in human health and aging. Front Microbiol 2016, 7:1204.

57. Hildebrand F, Moitinho-Silva L, Blasche S, Jahn MTT, Gossmann TI, Heuerta Cepas J, Hercog R, Luetge M, Bahram M, Pryszlak A, Alves RJ, Waszak SM, Zhu A, Ye L, Costea PI, Aalvink S, Belzer C, Forslund SK, Sunagawa S, Hentschel U, Merten C, Patil KR, Benes V, Bork P: Antibiotics-induced monodominance of a novel gut bacterial order. Gut 2019,

58. Dubourg G, Lagier JC, Armougom F, Robert C, Audoly G, Papazian L, Raoult D: High-level colonisation of the human gut by Verrucomicrobia following broad-spectrum antibiotic treatment. Int J Antimicrob Agents 2013, 41:149–155.

59. Shaw TD, Fairley DJ, Schneiders T, Pathiraja M, Hill RLR, Werner G, Elborn JS, McMullan R: The use of high-throughput sequencing to investigate an outbreak of glycopeptide-resistant Enterococcus faecium with a novel quinupristin-dalfopristin resistance mechanism. Eur J Clin Microbiol Infect Dis 2018, 37:959–967.

60. Lee RS, Goncalves da Silva A, Baines SL, Strachan J, Ballard S, Carter GP, Kwong JC, Schultz MB, Bulach DM, Seemann T, Stinear TP, Howden BP: The changing landscape of vancomycin-resistant Enterococcus faecium in Australia: a population-level genomic study. J Antimicrob Chemother 2018, 73:3268–3278.

61. Zhou X, Chlebowicz MA, Bathoorn E, Rosema S, Couto N, Lokate M, Arends JP, Friedrich AW, Rossen JWA: Elucidating vancomycin-resistant Enterococcus faecium outbreaks: the role of clonal spread and movement of mobile genetic elements. J Antimicrob Chemother 2018, 73:3259–3267.

62. Hayakawa M, Asahara T, Henzan N, Murakami H, Yamamoto H, Mukai N, Minami Y, Sugano M, Kubota N, Uegaki S, Kamoshida H, Sawamura A, Nomoto K, Gando S: Dramatic changes of the gut flora immediately after severe and sudden insults. Dig Dis Sci 2011, 56:2361–2365.

63. Ojima M, Motooka D, Shimizu K, Gotoh K, Shintani A, Yoshiya K, Nakamura S, Ogura H, Iida T, Shimazu T: Metagenomic analysis reveals dynamic changes of whole gut microbiota in the acute phase of intensive care unit patients. Dig Dis Sci 2016, 61:1628–1634.

64. Yeh A, Rogers MB, Firek B, Neal MD, Zuckerbraun BS, Morowitz MJ: Dysbiosis across multiple body sites in critically ill adult surgical patients. Shock 2016, 46:649–654.

65. Buelow E, Bello González TDJ, Fuentes S, de Steenhuijsen Piters WAA, Lahti L, Bayjanov JR, Majoor EAM, Braat JC, van Mourik MSM, Oostdijk EAN, Willems RJL, Bonten MJM, van Passel MWJ, Smidt H, van Schaik W: Comparative gut microbiota and resistome profiling of intensive care patients receiving selective digestive tract decontamination and healthy subjects. Microbiome 2017, 5:88.

66. Raymond F, Ouameur AA, Deraspe M, Iqbal N, Gingras H, Dridi B, Leprohon P, Plante PL, Giroux R, Berube E, Frenette J, Boudreau DK, Simard JL, Chabot I, Domingo MC, Trottier S, Boissinot M, Huletsky A, Roy PH, Ouellette M, Bergeron MG, Corbeil J: The initial state of the human gut microbiome determines its reshaping by antibiotics. ISME J 2016, 10:707–720.

67. Lankelma JM, van Vught LA, Belzer C, Schultz MJ, van der Poll T, de Vos WM, Wiersinga WJ: Critically ill patients demonstrate large interpersonal variation in intestinal microbiota dysregulation: a pilot study. Intensive Care Med 2017, 43:59–68.

68. Lamarche D, Johnstone J, Zytaruk N, Clarke F, Hand L, Loukov D, Szamosi JC, Rossi L, Schenck LP, Verschoor CP, McDonald E, Meade MO, Marshall JC, Bowdish DME, Karachi T, Heels-Ansdell D, Cook DJ, Surette MG: Microbial dysbiosis and mortality during mechanical ventilation: a prospective observational study. Respir Res 2018, 19:220.

69. Livanos AE, Snider EJ, Whittier S, Chong DH, Wang TC, Abrams JA, Freedberg DE: Rapid gastrointestinal loss of Clostridial Clusters IV and XIVa in the ICU associates with an expansion of gut pathogens. PLoS ONE 2018, 13:e0200322.

70. Thiemann S, Smit N, Strowig T: Antibiotics and the intestinal microbiome: Individual responses, resilience of the ecosystem, and the susceptibility to infections. Curr Top Microbiol Immunol 2016, 398:123–146.

71. Lange K, Buerger M, Stallmach A, Bruns T: Effects of antibiotics on gut microbiota. Dig Dis 2016, 34:260–268.

72. Modi SR, Collins JJ, Relman DA: Antibiotics and the gut microbiota. J Clin Invest 2014, 124:4212–4218.

73. Ianiro G, Tilg H, Gasbarrini A: Antibiotics as deep modulators of gut microbiota: between good and evil. Gut 2016, 65:1906–1915.

74. Bradley SJ, Wilson AL, Allen MC, Sher HA, Goldstone AH, Scott GM: The control of hyperendemic glycopeptide-resistant Enterococcus spp. on a haematology unit by changing antibiotic usage. J Antimicrob Chemother 1999, 43:261–266.

75. Wilcox MH: Gastrointestinal disorders and the critically ill. Clostridium difficile infection and pseudomembranous colitis. Best Pract Res Clin Gastroenterol 2003, 17:475–493.

76. Props R, Kerckhof FM, Rubbens P, De Vrieze J, Hernandez Sanabria E, Waegeman W, Monsieurs P, Hammes F, Boon N: Absolute quantification of microbial taxon abundances. ISME J 2017, 11:584–587.

77. Vandeputte D, Kathagen G, D’hoe K, Vieira-Silva S, Valles-Colomer M, Sabino J, Wang J, Tito RY, De Commer L, Darzi Y, Vermeire S, Falony G, Raes J: Quantitative microbiome profiling links gut community variation to microbial load. Nature 2017, 551:507–511.

78. Ruppé E, Ghozlane A, Tap J, Pons N, Alvarez AS, Maziers N, Cuesta T, Hernando-Amado S, Clares I, Martínez JL, Coque TM, Baquero F, Lanza VF, Máiz L, Goulenok T, de Lastours V, Amor N, Fantin B, Wieder I, Andremont A, van Schaik W, Rogers M, Zhang X, Willems RJL, de Brevern AG, Batto JM, Blottière HM, Léonard P, Léjard V, Letur A, Levenez F, Weiszer K, Haimet F, Doré J, Kennedy SP, Ehrlich SD: Prediction of the intestinal resistome by a three-dimensional structure-based method. Nat Microbiol 2019, 4:112–123.

79. Khelaifia S, Drancourt M: Susceptibility of archaea to antimicrobial agents: applications to clinical microbiology. Clin Microbiol Infect 2012, 18:841–848.

80. Kaleko M, Bristol JA, Hubert S, Parsley T, Widmer G, Tzipori S, Subramanian P, Hasan N, Koski P, Kokai-Kun J, Sliman J, Jones A, Connelly S: Development of SYN-004, an oral beta-lactamase treatment to protect the gut microbiome from antibiotic-mediated damage and prevent Clostridium difficile infection. Anaerobe 2016, 41:58–67.

81. de Gunzburg J, Ghozlane A, Ducher A, Le Chatelier E, Duval X, Ruppé E, Armand-Lefevre L, Sablier-Gallis F, Burdet C, Alavoine L, Chachaty E, Augustin V, Varastet M, Levenez F, Kennedy S, Pons N, Mentré F, Andremont A: Protection of the human gut microbiome from antibiotics. J Infect Dis 2018, 217:628–636.

82. Taur Y, Coyte K, Schluter J, Robilotti E, Figueroa C, Gjonbalaj M, Littmann ER, Ling L, Miller L, Gyaltshen Y, Fontana E, Morjaria S, Gyurkocza B, Perales MA, Castro-Malaspina H, Tamari R, Ponce D, Koehne G, Barker J, Jakubowski A, Papadopoulos E, Dahi P, Sauter C, Shaffer B, Young JW, Peled J, Meagher RC, Jenq RR, van den Brink MRM, Giralt SA, Pamer EG, Xavier JB: Reconstitution of the gut microbiota of antibiotictreated patients by autologous fecal microbiota transplant. Sci Transl Med 2018, 10

83. Bilinski J, Grzesiowski P, Sorensen N, Madry K, Muszynski J, Robak K, Wroblewska M, Dzieciatkowski T, Dulny G, Dwilewicz-Trojaczek J, Wiktor-Jedrzejczak W, Basak GW: Fecal microbiota transplantation in patients with blood disorders inhibits gut colonization with antibiotic-resistant bacteria: Results of a prospective, single-center study. Clin Infect Dis 2017, 65:364–370.

