## Additional File 2 for "Loss of microbial diversity and pathogen domination of the gut microbiota in critically ill patients"

| Patient ID | Age | Sex | Admission reason | Co-morbidities | Total length of stay in the hospital | Length of stay in the ICU | 90-day mortality | Number of samples for sequencing |
| --- | --- | --- | --- | --- | --- | --- | --- | --- |
| 2 | 64 | F | Subarachnoid haemorrhage | COPD^1^ | 29 | 15 | No | 12 |
| 4 | 75 | M | Aortic Aneurysm repair | COPD^1^ | Transfer from another hospital | 14 | Yes | 7 |
| 8 | 59 | M | Subarachnoid haemorrhage | Nil | 52 | 17 | No | 9 |
| 10 | 55 | M | Multiple trauma | COPD^1^ | 0 | 13 | No | 5 |
| 15 | 56 | M | CABG^2^ | NIDDM^3^; Hypertension, endocarditis, Aortic valve stenosis | 13 | 5 | No | 2 |
| 22 | 72 | M | Pneumonia | TB^4^; NIDDM^3^ | 70 | 6 | No | 2 |
| 24 | 59 | M | Drug induced hepatitis | Nil | 23 | 17 | Unknown | 6 |
| 25 | 46 | M | Intracerebral bleed | Hydrocephalus, alcoholic liver disease, intracerebral haemorrhage | 29 | 24 | Unknown | 16 |
| 29 | 80 | M | Subcapsular haematoma | Liver cancer | 16 | 18 | No | 5 |
| 31 | 43 | M | Subarachnoid haemorrhage | Hypertension; alcoholism | 18 | 19 | Yes | 5 |
| 35 | 59 | M | Lung transplant | COPD^1^ | 41 | 14 | No | 9 |
| 36 | 30 | M | Multiple trauma |  | 50 | 20 | No | 8 |
| 37 | 47 | M | Multiple trauma | Depression | 62 | 27 | No | 9 |
| 38 | 47 | M | Insertion of left ventricular assist device | NIDDM^3^, essential hypertension | 133 | 60 | No | 23 |
| 41 | 41 | M | Oesophagostomy | Oesophageal adenocarcinoma | 144 | 45 | No | 10 |
| 45 | 63 | M | Multiple trauma |  | 51 | 26 | No | 11 |
| 46 | 25 | M | Bacterial pneumonia | Nil | 44 | 37 | Yes | 12 |
| 47 | 46 | M | Acute subdural haematoma | Hepatitis C and schizophrenia | Unknown | 20 | No | 7 |
| 49 | 65 | F | Intracerebral haematoma | Breast cancer | 28 | 11 | No | 6 |
| 51 | 78 | M | ST-elevation myocardial infarction | Nil | 37 | 27 | No | 6 |
| 52 | 54 | F | Aortic surgery | Suspected endocarditis | 59 | 20 | No | 5 |
| 53 | 40 | F | Anaemia | End stage renal disease | 94 | 42 | No | 21 |
| 54 | 66 | M | Alcohol withdrawal syndrome | Epilepsy | 47 | 32 | Yes | 8 |
| 55 | 66 | F | Subdural haemorrhage | NIDDM^3^ | 56 | 55 | No | 19 |
| 57 | 84 | M | Cardiac arrest | Hypertension; Cardiomyopathy | 108 | 14 | No | 5 |
| 59 | 77 | M | Subdural haematoma | Hyperlipidaemia; Hypertension | 56 | 28 | No | 6 |

^1^Chronic Obstructive Pulmonary Disease; ^2^ Coronary Artery Bypass Grafting; ^3^Non-Insulin Dependent Diabetes Mellitus; ^4^Tuberculosis
