## Supplementary figures and images for "Loss of microbial diversity and pathogen domination of the gut microbiota in critically ill patients"

### Additional File 5

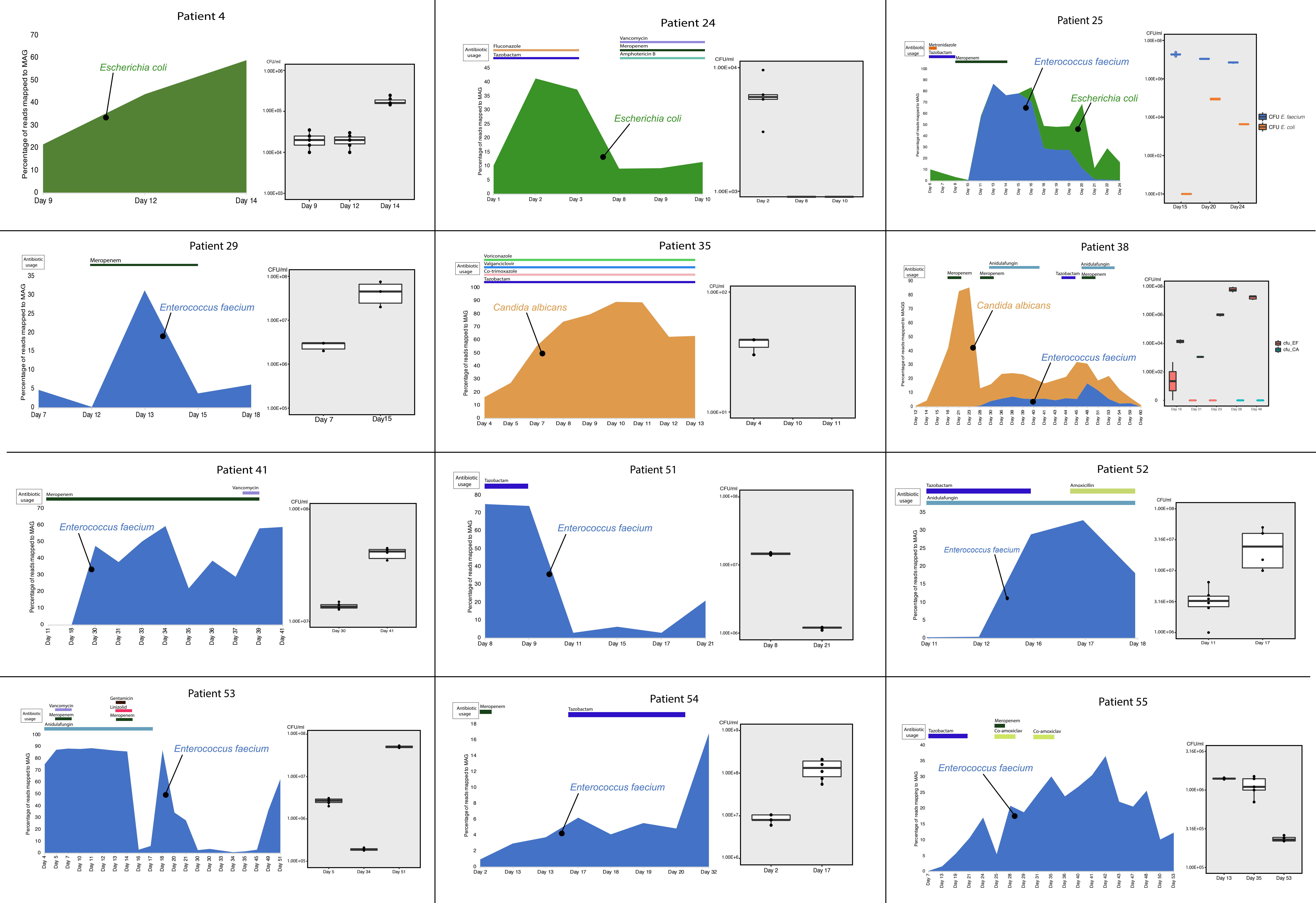
