## Additional File 6 for "Loss of microbial diversity and pathogen domination of the gut microbiota in critically ill patients"

| Patient Name | MAG identified | Reference Size | Completeness | Contamination | MLST |
| --- | --- | --- | --- | --- | --- |
| Patient25 | *E. faecium* | 2,987,425 | 99.63 | 0.87 | 787 |
| Patient29 | *E. faecium* | 2,769,265 | 98.88 | 0.06 | - |
| Patient38 | *E. faecium* | 2,689,224 | 99.63 | 0 | 262 |
| Patient41 | *E. faecium* | 2,944,903 | 99.63 | 0.5 | 80 |
| Patient51 | *E. faecium* | 2,908,317 | 99.63 | 0.56 | 80 |
| Patient52 | *E. faecium* | 2,843,541 | 98.88 | 0 | 80 |
| Patient53 | *E. faecium* | 2,837,393 | 99.25 | 0.12 | 80 |
| Patient54 | *E. faecium* | 2,421,346 | 99.3 | 0.06 | 80 |
| Patient55 | *E. faecium* | 2,979,337 | 99.44 | 0.56 | 80 |
| Patient4 | *E. coli* | 4,015,777 | 76.47 | 2.29 | - |
| Patient25 | *E. coli* | 4,970,123 | 98.65 | 1.24 | 315 |
| Patient36 | *E. coli* | 4,288,275 | 97.57 | 0.11 | 538 |
| Patient24 | *E. coli* | 4,646,804 | 83.17 | 2.09 | - |
| Patient35 | *C. albicans* | 16116472 | 83.1325 | 14.1 | - |
| Patient38 | *C. albicans* | 15296317 | 76 | 7.2 | - |
| Patient 31 | *P. mirobilis* | 3,819,203 | 100 | 0 |  |
| Patient2 | *K. pneumoniae* | 4,782,063 | 87.58 | 8.9 |  |
| Patient38 | *Enterobacter sp.* | 4,884,100 | 99.9 | 0.4 |  |
