## Additional File 8 for "Loss of microbial diversity and pathogen domination of the gut microbiota in critically ill patients"

Supplementary Table 7

Identification of antibiotic resistance genes in the metagenome assembled genomes

***Enterococcus faecium***

| Patient Name | Pathogen MAG | | Aminoglycosides | Macrolides and Streptogramin B | Glycopeptides | | | |
| --- | --- | --- | --- | --- | --- | --- | --- | --- |
| Antibotic resistance genes | | | *Aph*(3) | *Msr*C | *Van*A | *Van*H | *Van*X | *Van*Z |
| Patient 25 | | *Enterococcus faecium* |  | 100 | 100 | 100 | 100 |  |
| Patient 29 | |  |  | 100 |  |  |  |  |
| Patient 38 | |  |  | 100 |  |  |  |  |
| Patient 41 | |  |  | 100 |  |  |  |  |
| Patient 51 | |  |  | 100 |  |  |  |  |
| Patient 52 | |  |  | 100 |  |  |  |  |
| Patient 53 | |  | 100 | 100 | 100 | 100 | 100 | 100 |
| Patient 54 | |  |  | 100 |  |  |  |  |
| Patient 55 | |  |  | 100 |  |  |  |  |

***Escherichia coli***

| Patient Name | Pathogen MAG | Aminoglycosides | | Macrolides and Streptogramin B | | | Betalactam | |
| --- | --- | --- | --- | --- | --- | --- | --- | --- |
| Antibiotic resistance genes | | *Aac*(3) | *Ant*(3) | *mef*B | *mph*A | *mdh*A | *bla*CTX | *bla*TEM |
| Patient 4 | *Escherichia coli* | 99.9 |  | 99.6 | 99.7 | 99.8 |  |  |
| Patient 25 |  | 99.8 | 99.3 |  | 99.6 | 98.2 | 100 | 100 |
| Patient 36 |  |  |  |  |  | 98 |  |  |
| Patient 24 |  |  |  |  |  | 98.3 |  |  |

**Other MAGS**

| Patient Name | Pathogen MAG | Beta-lactams | | Quinolones | | Antifungal agents |
| --- | --- | --- | --- | --- | --- | --- |
| Antibiotic resistance genes | | *bla*SHV | *bla*ACT | *oqx*A | *oqx*B | Fluconazole (ERG11) |
| Patient 2 | *Klebsiella pneumoniae* | 99.65 |  | 100 | 99.3 |  |
| Patient 38 | *Enterobacter sp.* |  | 99.83 |  |  |  |
| Patient 38 | *Candida albicans* |  |  |  |  | 99.43 |
| Patient 35 | *Candida albicans* |  |  |  |  | 99.37 |
